# Polymorphonuclear neutrophils modulate the responses of human immune cells to vaccines in an *in vitro* blood cell culture system

**DOI:** 10.64898/2026.06.15.732289

**Authors:** Shuran Gong, Harshad P. Patil, Jacqueline De Vries-Idema, Martin Beukema, Anke Huckriede

## Abstract

Vaccine-induced immune responses are the result of an intricate interplay between different cell populations of the innate and adaptive immune system, which is so far only partly understood. In particular, the role of polymorphonuclear neutrophils (PMNs) has long been neglected. Here, we studied the effects of a whole inactivated virus influenza vaccine (WIV) in an *in vitro* system consisting of freshly isolated human PMNs alone or PMNs combined with autologous peripheral blood mononuclear cells (PBMCs). Isolated PMNs showed minimal responses to the vaccine with respect to apoptosis, gene expression, cytokine production, and reactive oxygen species production. However, in WIV-stimulated PMN/PBMC co-cultures, PMNs particularly enhanced monocyte dynamics, CD14^-^CD11c^+^ cell activation, effector T cell differentiation, and B cell antibody production. On the other hand, PMNs decreased T follicular helper cell frequencies. Without vaccine stimulation, PMN presence resulted in enhanced levels of baseline inflammatory cytokines in PMN/PBMC co-cultures. However, with vaccine stimulation, PMNs dampened the vaccine-induced cytokine secretion of PBMCs. These findings reveal PMNs as regulators of vaccine responses whose effects depend on crosstalk with other immune cells, balancing pro-inflammatory and adaptive immune activation.

**Author summary:** Polymorphonuclear neutrophils (PMNs) are essential and predominant cells of the human innate immune system. Growing evidence implicates that PMNs are involved in vaccine-induced immune activation, but their exact role is so far poorly defined. In our study, human PMNs were tested alone to observe their response to whole inactivated virus influenza vaccine (WIV), or combined with autologous peripheral blood mononuclear cells (PBMCs) to investigate how their presence influences vaccine responses of various cell populations within PBMCs. Our results show that WIV had little direct effect on isolated PMNs. However, when PMNs were combined with other immune cells, PMNs acted as crucial regulators: they enhanced the activity of innate immune cells, regulated the responses to the vaccine of T and B cells, and helped control the overall level of inflammation. Our study forms the groundwork for a more comprehensive understanding of human immune cell interactions under vaccine stimulation.

## Introduction

Vaccines are highly instrumental in fighting bacterial and viral infections through the induction of pathogen-specific immune responses [1, 2]. These immune responses are the result of an intricate interplay between cells of the innate and adaptive immune system. However, studies on the effect of vaccines on immune cells have largely focused on dendritic cells as representatives of the innate immune system and T and B cells of the adaptive immune system [3–6]. Recently, evidence has been accumulating for a role of polymorphonuclear neutrophils (PMNs) in vaccine-induced immune responses. As first responders to infection or vaccination, PMNs rapidly migrate to sites of antigen exposure, where they are among the earliest cells to encounter pathogens and initiate immune activation [7]. In mice and macaques, various types of vaccines were shown to recruit PMNs to the site of immunization, where they get activated and help in shaping an inflammatory environment, attracting other immune cells, and/or trafficking antigens to draining lymph nodes [8–14]. Several studies demonstrate that depletion of PMNs around the time of vaccination impairs CD4^+^ T cell induction, antibody affinity, and antibody class switching, and might even abrogate protective responses completely [12, 15–17]. However, all these results were obtained from animal models, which are known to differ from humans regarding PMN frequency and the expression of relevant PMN (surface) markers [18, 19]. Information on the potential role of PMNs in human vaccination is still very limited. Therefore, this *in vitro* study aimed to determine how human PMNs respond to vaccines and how they influence vaccine-induced activation of other immune cells.

To investigate the effects of vaccines on human neutrophils and the impact of these effects on other immune cells, we used a whole inactivated virus influenza vaccine (WIV) as a model vaccine. WIV is unable to replicate but retains the structure and the single-stranded RNA of the native influenza virus and activates immune cells via engagement of Toll-like receptor 7 [20, 21]. To determine whether PMNs can directly respond to the vaccine, we treated isolated PMNs with WIV, and investigated their responses, focusing on apoptosis, expression of activation markers, and the production of cytokines and reactive oxygen species (ROS). Subsequently, to reveal the role of PMNs in vaccine-evoked responses, we co-cultured human-derived autologous PMNs and PBMCs at different ratios and assessed vaccine-induced responses of innate and adaptive immune cells. Our results indicate that PMNs alone show limited responses to WIV. However, when co-cultured with PBMCs, they affect the responses of other immune cell populations. These observations imply that PMN responses should be taken into account when designing improved vaccines and adjuvants.

## Methods

### Vaccines and positive stimulants

The NIBRG-23 virus, a recombinant reassortant of A/turkey/Turkey/1/2005 (H5N1) and A/Puerto Rico/8/34 (H1N1), originated from the National Institute for Biological Standards and Control (Potters Bar, United Kingdom) and was propagated on Madin-Darby canine kidney (MDCK) cells. WIV was prepared using this strain, following previously described protocols [5]. The TLR4 ligand lipopolysaccharide (LPS; standard grade, from *Escherichia coli* K12, Invivogen, Toulouse, France), the TLR7/8 ligand resiquimod (R848; Invivogen, Toulouse, France), and the cytokine interleukin 2 (IL-2; PeproTech, London, UK) were aliquoted in concentrated solution and preserved at -20°C.

### Cell isolation

Autologous human PMNs and PBMCs were isolated from buffy coats of healthy volunteers, obtained from the Dutch Blood Bank (Sanquin, Nijmegen, NL). PMNs and mononuclear cells were separated using density gradient centrifugation (800xg, 30min) with Ficoll Histopaque, and PBMCs were collected as described previously [3]. For PMN isolation, all layers above the PMN-erythrocyte layer were removed from the centrifuged gradient. The PMN-erythrocyte layer was transferred to a new tube, and the erythrocytes were lysed using ice-cold ACK lysing buffer (Gibco Life Technologies, NY, USA) at a ratio of 2.5:1 (vol:vol ACK buffer to granulocyte-erythrocyte mixture). The tube was gently inverted every 30s during lysis. Once the solution turned black and viscosity dropped, Hank’s balanced solution (HBSS; Ca/Mg ^-/-^, Gibco Life Technologies, Paisley, UK) with 5% fetal bovine serum (FBS; Life Science Production, Bedfordshire, UK) was added to a final volume of 50 mL, followed by centrifugation at 200xg for 3 mins. The supernatant was discarded, and the lysis step was repeated if erythrocytes were still visible. Subsequently, PMNs were suspended in ice-cold complete medium (RPMI 1640, HEPES (Gibco Life Technologies, Paisley, UK) supplemented with 10% heat-inactivated FBS, and 100 U/mL L-glutamine (both Gibco Life Technologies, Paisley, UK)), and live cells were counted using Trypan Blue solution (Merck Life Science, Gillingham, UK). To prevent pre-activation, all buffers and media were pre-cooled on ice.

### Culture and stimulation of human PMNs and PBMC *in vitro*

Isolated PMNs were cultured alone (1x10^6^ cells per mL per well) or were co-cultured with autologous PBMCs at PMN:PBMC ratios of 2:1, 1:1, 0.3:1, or 0:1 in 1 mL volumes in 24-well plates. The PBMC number was maintained at 0.5x10^6^ cells per well. Complete medium was used for all experiments. WIV (at the indicated final concentrations), LPS (0.1 µg/mL) or R848 + IL-2 (respectively 1 µg/mL, InvivoGen, Toulouse, France and 10 ng/mL, PeproTech, London, UK) was added to each (mixed) cell culture. Cells were harvested on days 1, 3, and 5 for flow cytometry. Supernatants were collected for quantification of total IgG by ELISA and cytokines/chemokines by multiplex bead-based immunoassay *(see below)*.

### Multi-color flow cytometry

On day 1, cells were harvested into a 96-DeepWell plate (Thermo Fisher Scientific Co., Carlsbad, CA, USA) and incubated with antibodies to assess innate immune responses (LIVE/DEAD™ Fixable Blue, CD11c, CD14, CD15, CD16, HLA-DR, CD86, CD80, CD40; see supplementary Table 1 for details on antibodies used) for 30 minutes in FACS buffer (1x Dulbecco’s phosphate-buffered saline (DPBS; Gibco) supplemented with 2% FBS) at 4°C. After washing with FACS buffer, cells were fixed with 4% paraformaldehyde (PFA). On day 3 or day 5, prior to sample collection, brefeldin A (1:1000, BioLegend, Fell, DE) was added to the medium for 6 h. Subsequently, cells were harvested, washed, and incubated with antibodies targeting surface antigens of adaptive immune cells (CD3, CD4, CD8, CXCR5, ICOS, CD45RA, CCR7, CD19, CD20, CD27, CD38; see supplementary Table 1 for further details) for 30 mins at 4 ℃. Following staining, the cells were fixed and permeabilized using the BD Cytofix/Cytoperm™ Fixation/Permeabilization Kit (BD Biosciences, San Diego, USA). The stained and washed cells were recorded using a Cytek Aurora flow cytometer (Cytek Biosciences, Amsterdam, The Netherlands). The unmixing plots of each sample were extracted and analyzed with Spectroflo software (Cytek Biosciences) and Kaluza software (Beckman Coulter, Woerden, NL). Fluorescence minus one (FMO) controls were applied to determine the positive gates; gating strategies are shown in *Supplementary Figure 1*.

### Cytokine quantification by LEGENDplex ™ bead-based immunoassays

Supernatants were collected from PMN cultures after 20 h of *in vitro* stimulation and from PMN:PBMC co-cultures (ratio 2:1) and PBMC cultures after 24 hours of stimulation. Cytokine levels were measured using a LEGENDplex™ HU Essential Immune Responses Panel (13-plex) kit (Ref num: 740930 Lot: B338757, BioLegend, San Diego, CA, USA) according to the manufacturer’s instructions. Data acquisition and analysis were performed using LEGENDplex™ Data Analysis Software (BioLegend). Cytokine concentrations of the samples were determined from the standard curve using a 5-parameter logistic curve-fitting method.

### Annexin V-PI double staining with flow cytometry

To assess cell apoptosis, Annexin V-FITC and propidium iodide (PI) double staining was performed as previously described [22]. Briefly, cells were harvested and washed twice with cold phosphate-buffered saline (PBS). Cell pellets were resuspended in 2 mL of 1x binding buffer (10 mM HEPES, 140 mM NaCl, 2.5 mM CaCl_2_, pH 7.4) and centrifuged for 6 min at 375 x g. Annexin V-FITC (2.5 µL) was added to the cell suspension, and the mixture was gently vortexed. The cells were incubated for 15 minutes at RT in the dark. After incubation, cells were washed once with binding buffer. The PI (3 µL) was added 3 mins before sample analysis. Stained samples were analyzed using a flow cytometer (Cytek Aurora, Cytek Biosciences, Amsterdam, NL). Data acquisition and analysis were performed using Kaluza software (Beckman Coulter, Woerden, NL). Single stainings with mixed cells were performed for unmixing channel extraction and gate setting.

### Measurement of intracellular reactive oxygen species (ROS) using CM-H₂DCFDA

Intracellular ROS levels were measured using the fluorescent probe CM-H₂DCFDA (5-(and-6)-chloromethyl-2’,7’-dichlorodihydrofluorescein diacetate, acetylated form; molecular weight: 577.8 g/mol) [23]. A stock solution (2 mM) was prepared by dissolving 50 µg of CM-H₂DCFDA in 43.25 µL of dimethyl sulfoxide (DMSO). The working solution (5 µM) was prepared in serum-free medium to prevent extracellular cleavage of acetyl groups by serum esterases. Cells seeded in a 24-well plate were washed, and 300 µL of serum-free medium containing 2.5 µM CM-H₂DCFDA (diluted 1:500 from the stock) was added per well. After incubation for 30 minutes at 37°C, the dye-containing medium was removed, and cells were incubated for an additional 30 minutes in normal growth medium to facilitate complete deacetylation and oxidation of the probe. Following incubation, cells were washed with 1x PBS, centrifuged at 2,000 rpm for 5 minutes, and resuspended in 200 µL PBS. Samples were kept on ice and protected from light until analysis. Fluorescence was measured using a Guava easyCyte Flow Cytometer (Luminex, Austin, Texas, USA) with excitation at 492–495 nm and emission detection at 517–527 nm.

### Determination of total IgG

Cryopreserved supernatants of vaccine-stimulated PBMC cultures, with and without PMNs, collected on day 5, were thawed and diluted 2-10 times. Total IgG was determined by ELISA as described previously [5]. Absorbance was quantified at a wavelength of 450 nm using a microtiter plate reader. The IgG levels in each sample were determined by regression analysis using a titration curve of standard IgG.

### Data analysis

Statistical analyses were conducted using GraphPad Prism software (version 10.4.2, GraphPad Software, San Diego, CA, USA). To evaluate significant differences among multiple related groups, the nonparametric Friedman test was applied.

## Results

### Vaccine-stimulated PMNs exhibit minimal activation but slightly altered apoptosis dynamics

To gain insight into the potential role of PMNs in vaccination, we first investigated responses of freshly isolated human PMNs to stimulation with WIV, focusing on apoptosis, antigen presentation capability, reactive oxygen species levels, and the surface expression of activation markers.

Mature PMNs are known to have a short lifespan *in vitro* [24]. After culturing PMNs for 20h, most cells had indeed gone into early or late apoptosis, irrespective of the treatment. However, we observed slightly elevated percentages of live PMNs and slightly decreased percentages of early and late apoptotic PMNs upon stimulation with WIV (10, 5, 2.5, 1.25, and 0.625 µg/mL final concentration) (Fig. 1A-C). However, these differences did not reach statistical significance. The proportion of necrotic cells was very low under all conditions (Fig. 1D).

**Fig 1.**
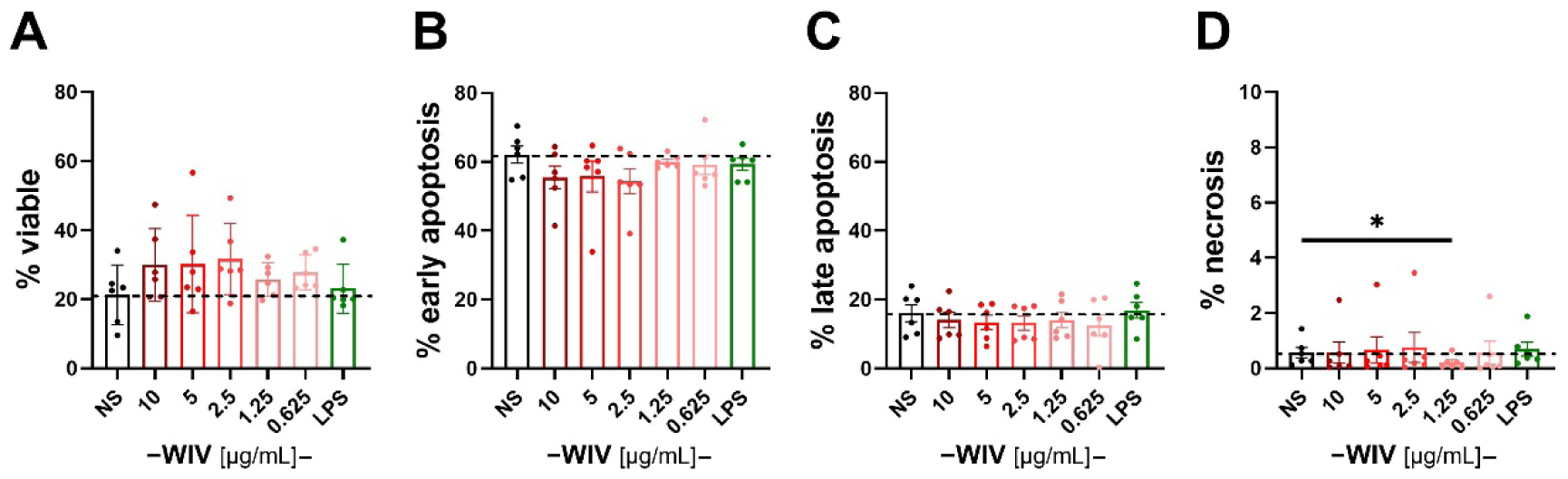
Evaluation of vaccine effects on cell death of cultured PMNs. PMNs, freshly isolated from buffy coats (n=6), were cultured without stimulus (non-stimulated (NS)), stimulated with WIV (at the indicated concentrations [µg/mL]), or LPS (0.1 µg/mL) for 20h and subsequently stained with Annexin V and PI. Flow cytometry was employed to measure the percentages of live cells (A), early apoptotic cells (B), late apoptotic cells (C), and necrotic cells (D). Dashed lines indicate the average percentages of the respective cells in NS cultures for comparison. Statistically significant differences between groups were determined using the paired Friedman test, with *p < 0.05 indicating significance.

We next assessed the activation status of PMNs after exposure to stimuli. Neither stimulation with WIV, nor with LPS used as positive control, enhanced the surface expression of the antigen presentation-related activation markers HLA-DR, CD86, and CD80 over that of unstimulated PMNs (Fig. 2 A-C). Similarly, the vaccine had no effect on ROS production (Fig. 2 D) and cytokine secretion, except for a slight increase in MCP-1 secretion by WIV-stimulated PMNs (Fig. 2 E-I). In contrast, LPS stimulated the production of ROS (Fig. 2D) and the secretion of IL-6 and IL-8, and to a lesser extent TNFα and CXCL-10, thus confirming the general responsiveness of the PMNs (Fig 2 E, F).

**Fig. 2.**
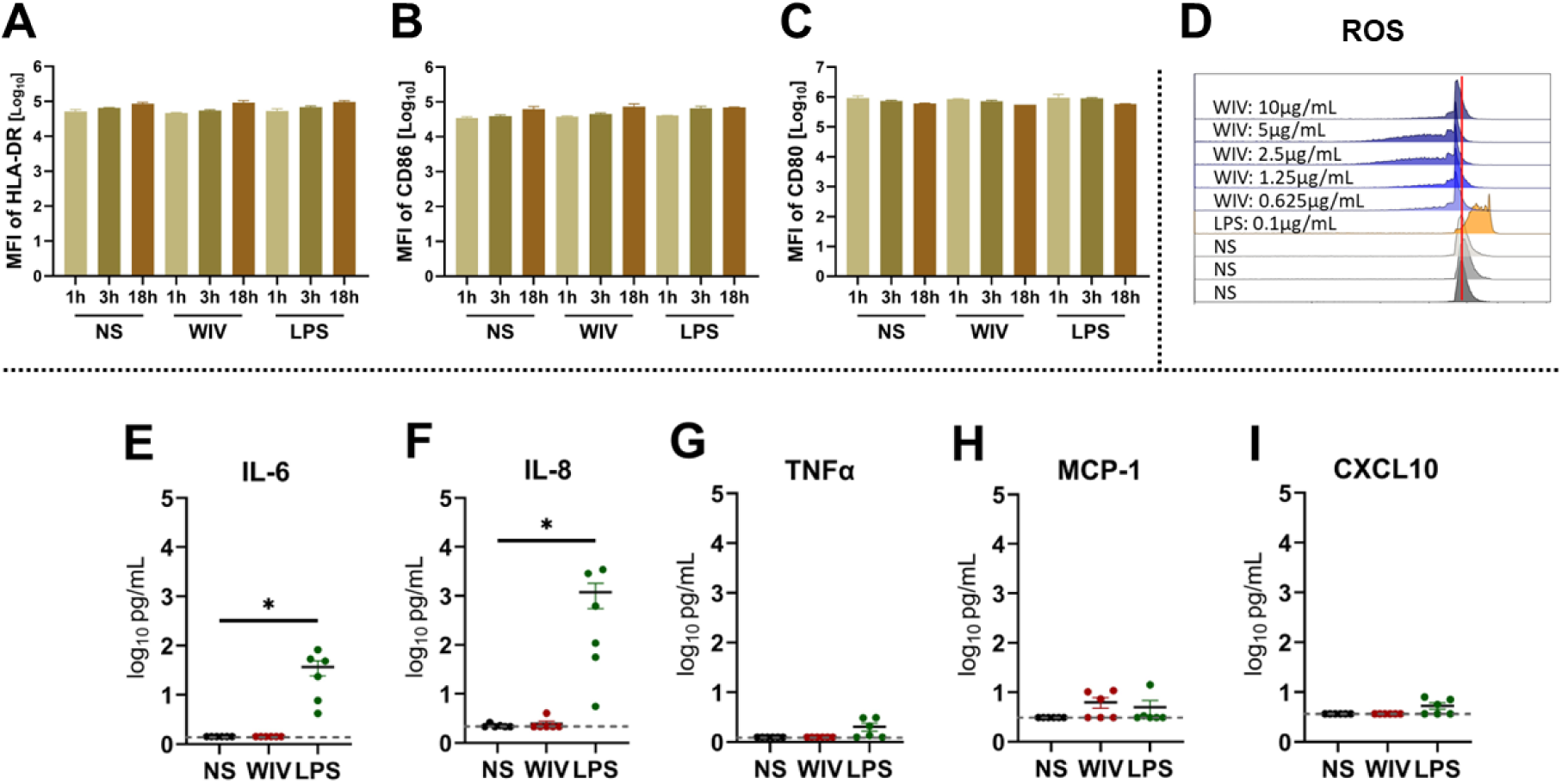
Activation of PMNs in response to WIV and LPS. Freshly isolated PMNs were left untreated (NS) or cultured in the presence of WIV (10 µg/mL) or LPS (0.1 µg/mL). The surface expression of HLA-DR (A), CD86 (B), and CD80 (C) was assessed by flow cytometry after 1, 3, and 18 h of incubation of the cells with WIV or LPS (n=2). D: Reactive oxygen species (ROS) generation was assessed after 4 h of stimulation of PMNs with WIV (0.625, 1.25, 2.5, 5, or 10 µg/mL) or LPS (0.1 µg/mL) by DCFH-DA staining followed by flow cytometry (n=2). E-I: The supernatants of the stimulated cells were harvested after 20 h and cytokine and chemokine levels were determined using the LEGENDplex multiplex bead-based immunoassay (n=6). Statistically significant differences between groups were assessed using the paired Friedman test, with *p < 0.05 denoting a significant difference.

Taken together, our results show that PMNs cultured *in vitro* exhibited little response to WIV influenza vaccine and might require further cytokines or additional immune cells to get fully activated. Consequently, we co-cultured PMNs with other immune cells to further assess potential mutual interactions.

### PMNs modulate WIV-induced monocyte subset changes and CD14^-^CD11c^+^cell activation in PBMC co-cultures

To elucidate possible interactions of PMNs with innate and adaptive immune cells present in PBMCs and vice versa, PMNs were co-cultured with autologous PBMCs at varying ratios and stimulated with WIV or LPS. Cells were collected at different time points to assess changes in cell numbers, subtypes, and activation levels using flow cytometry.

PMNs, when stimulated with GM-CSF or IFNγ, or co-cultured with T cells, can acquire antigen-presenting capabilities, despite exhibiting minimal or negligible expression of MHC-II and costimulatory molecules when cultured alone [25, 26]. However, our research indicated that the expression levels of HLA-DR, CD86, and CD80 on PMNs co-cultured with PBMCs at different ratios remained unchanged (Fig. S2 A, E, and I). Neither stimulation with WIV (Fig. S2 B, F, J) nor with LPS (Fig. S2 C, G, K) or a combination of IL2 and the TLR7/8 ligand R848 (Fig. S2 D, H, L) changed this result. Interestingly, we found that under WIV stimulation, PMNs co-cultured with PBMCs maintained the surface expression of CD16, which was strongly diminished in unstimulated cultures (Fig. S3 A and B). CD16, the Fcγ receptor IIIa, is involved in antibody-dependent cellular cytotoxicity (ADCC) and phagocytosis [27, 28]. High expression typically indicates mature, activated PMNs capable of enhanced immune responses [29–31].

To assess the effect of PMNs on the responses of other innate immune cells, we measured the proportions of monocyte subsets in PBMCs in the absence and presence of PMNs. In line with earlier reports [55], we observed that in freshly isolated PBMCs classical monocytes (CMs; CD14^hi^CD16^−^) accounted for the majority of monocytes (96.33%±0.63), followed by intermediate monocytes (IMs; CD14^hi^CD16^+^; 1.27%±1.13); non-classical monocytes (NCMs; CD14^+^CD16^+^) were not observed (results not shown). However, after 24 hours of culture, as employed in the following experiments, IMs became the predominant subtype (Fig. 3 B). In the absence of stimuli, the addition of PMNs did not alter the relative frequencies of the different monocyte subsets (Fig. 3 B). Stimulation of PBMCs with WIV increased the proportions of CMs, NCMs, and CD14^-^CD16^-^ cells (compared to those in non-stimulated co-cultures) while the proportion of intermediate monocytes decreased (Fig. 3 C, 0:1 condition). In PMN/PBMC co-cultures, this effect of WIV stimulation was sustained but less pronounced as the percentages of IM increased when more PMNs were present. A strong increase of CMs at the expense of IMs was observed for cultures stimulated with LPS or R848/IL-2 (Fig. 3 D, E), irrespective of PMNs being present or absent in the cultures.

**Fig 3.**
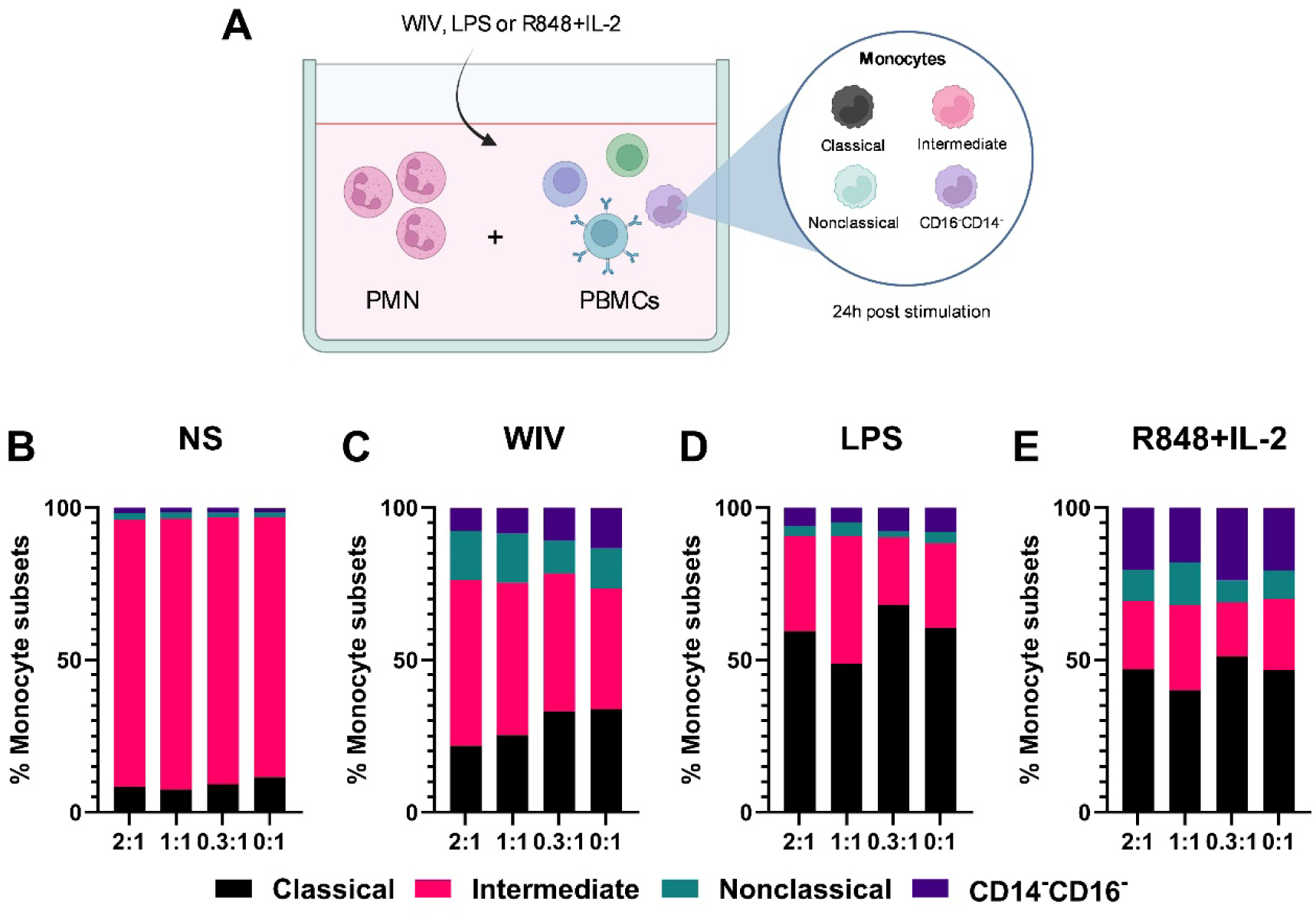
Effects of vaccine and TLR ligands on monocyte subsets. (A) Experimental setup: autologous human PMNs and PBMCs were co-cultured in the absence and presence of stimuli and the relative frequencies of classical (CD14^hi^CD16^−^), intermediate (CD14^hi^CD16^+^), and nonclassical (CD14^+^CD16^+^) monocytes and CD14^-^CD16^-^ cells were determined by flow cytometry. B – E: Cultures containing PMNs and PBMCs at the indicated ratios were left untreated (B) or stimulated with WIV (C), LPS (D), or R848+IL2 (E) for 24 h. Monocyte subsets in PBMCs were determined by flow cytometry based on CD14 and CD16 expression.

In the same cultures, we also assessed the quantity and activation status of CD14^-^CD11c^+^ cells, a population consisting mainly of classical dendritic cells [32]. CD14^-^CD11c^+^ cells possess a robust capacity to acquire antigens, thereby promoting T cell stimulation [33]. The expression of immunostimulatory molecules (MHC class II, CD86, CD80, and CD40) serves as indication of antigen-presenting cell maturation or activation [33, 34]. In the absence of stimuli (NS), co-culturing of PBMCs with PMNs for 24h did not significantly change the frequency of CD14^-^ CD11c^+^ cells (Fig. S4 A). In cultures stimulated with WIV, LPS, or R848+IL-2 (Fig. S4 B - D), the CD14^-^CD11c^+^ cell frequency increased for most donors. However, these changes were not affected by the number of PMNs present in the cultures.

Regarding the activation status of the antigen-presenting cells, PMNs had inconsistent effects (Fig. 4). In the absence of stimuli (NS), increasing numbers of PMNs enhanced the expression of HLA-DR and CD86 on CD14^-^CD11c^+^ cells of most donors, although these effects were not significant. In WIV-stimulated cultures, increasing PMN numbers resulted in enhanced expression of CD86, and a trend towards increased expression of CD80 on CD14^-^CD11c^+^ cells. In contrast, in LPS- and R848+IL-2-stimulated cultures, effects were minimal or inconsistent.

**Fig 4.**
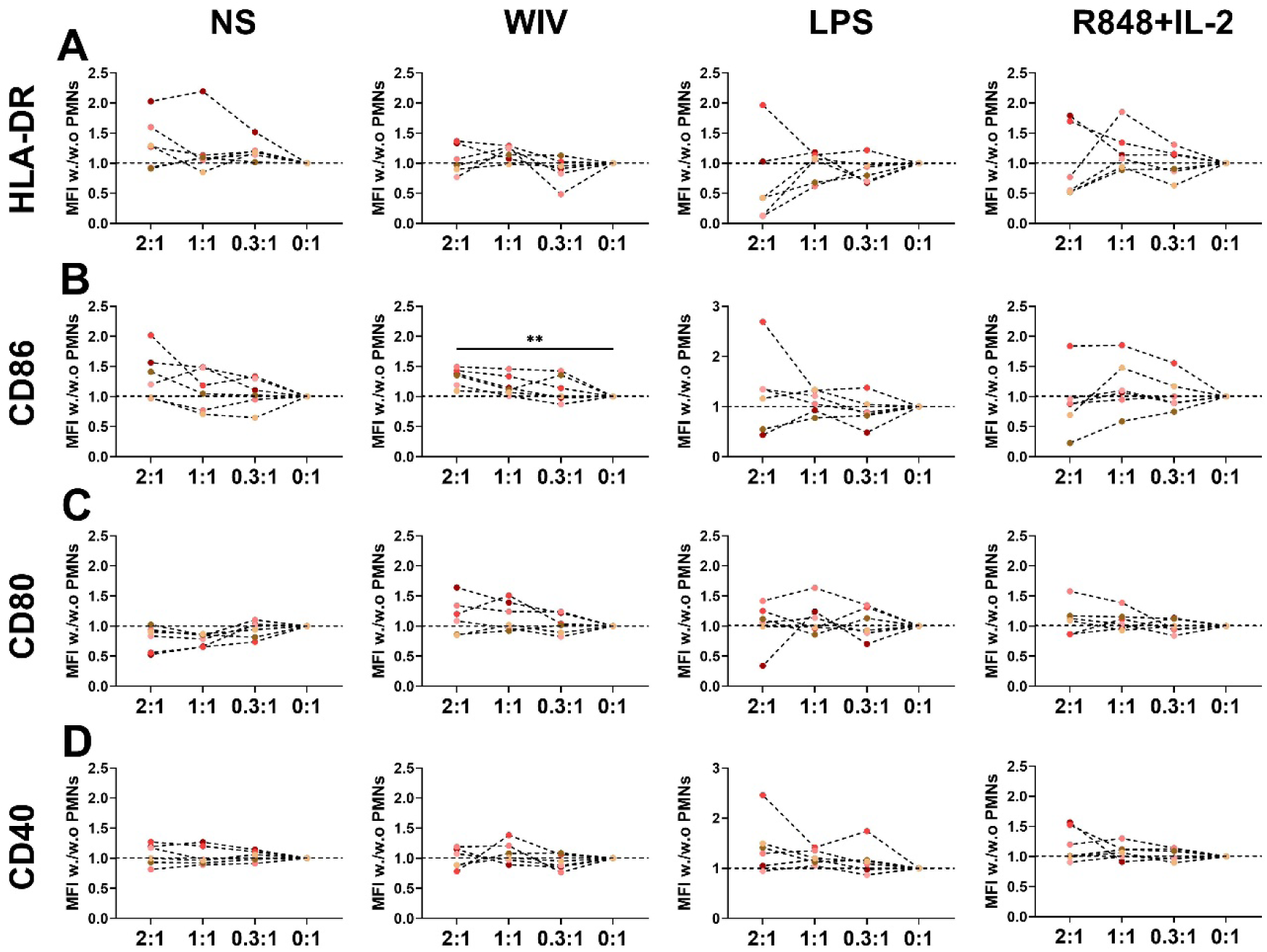
Activation status of CD14^-^CD11c^+^ cells in PMN/PBMC co-cultures. PBMCs alone (0:1) or supplemented with PMNs at the indicated ratios were stimulated for 24 h with WIV, LPS, or R848+IL-2, or left untreated. The mean fluorescence intensities (MFI) of the activation markers HLA-DR (A), CD86 (B), CD80 (C), and CD40 (D) on CD14^-^CD11c^+^ cells were determined by flow cytometry. Data is presented as the fold change of activation marker expression MFI in the presence of PMNs (2:1, 1:1, 0.3:1 ratios) divided by the MFI in the absence of PMNs (0:1 ratio) for the respective stimulation. Statistically significant differences between groups were assessed using the paired Friedman test.

In summary, after 24 h of co-culturing PMNs with PBMCs, there were no significant changes in activation marker expression on PMNs nor in the proportions of monocyte subsets or the number of CD14^-^CD11c^+^ cells in PBMCs. Nevertheless, PMNs modulated WIV-induced changes in monocyte subset frequencies and slightly enhanced the expression of some activation markers on non-stimulated as well as vaccine-stimulated CD14^-^CD11c^+^ cells.

### PMNs modulate T cell responses in PBMC co-cultures by promoting T effector cell differentiation while suppressing T follicular helper cells

We next studied the effects of PMNs on adaptive immune cells and their responses to vaccines by stimulating human PMN/PBMC co-cultures with WIV for 3 or 5 days. Irrespective of the number of PMNs added and the type of vaccine present, PMNs had little impact on the frequencies of B and T cells among the lymphoid cells or the proportions of CD4^+^ and CD8^+^ cells among CD3^+^ T cells (Fig. S5).

Zooming into T cell subsets, we observed that, after 3 and 5 days of culture without stimulation, PMNs slightly enhanced the differentiation of CD4^+^ and CD8^+^ naïve T cells (T_N_) into effector memory T cells (T_EM_) in a ratio-dependent manner but barely had an effect on central memory T cells (T_CM_) (Fig. 5 B-E). Similar effects were seen upon stimulation with WIV, butfor CD4 + T cells, the proportions of T_N_ were consistently higher and those of T_CM_ were consistently lower than in non-stimulated cultures at the same PMN:PBMC ratio. For CD8+ T cells, the addition of PMNs to WIV-stimulated cultures slightly elevated the proportion of terminally differentiated T cells (T_EMRA_) (Fig. 5 D, E).

**Fig 5.**
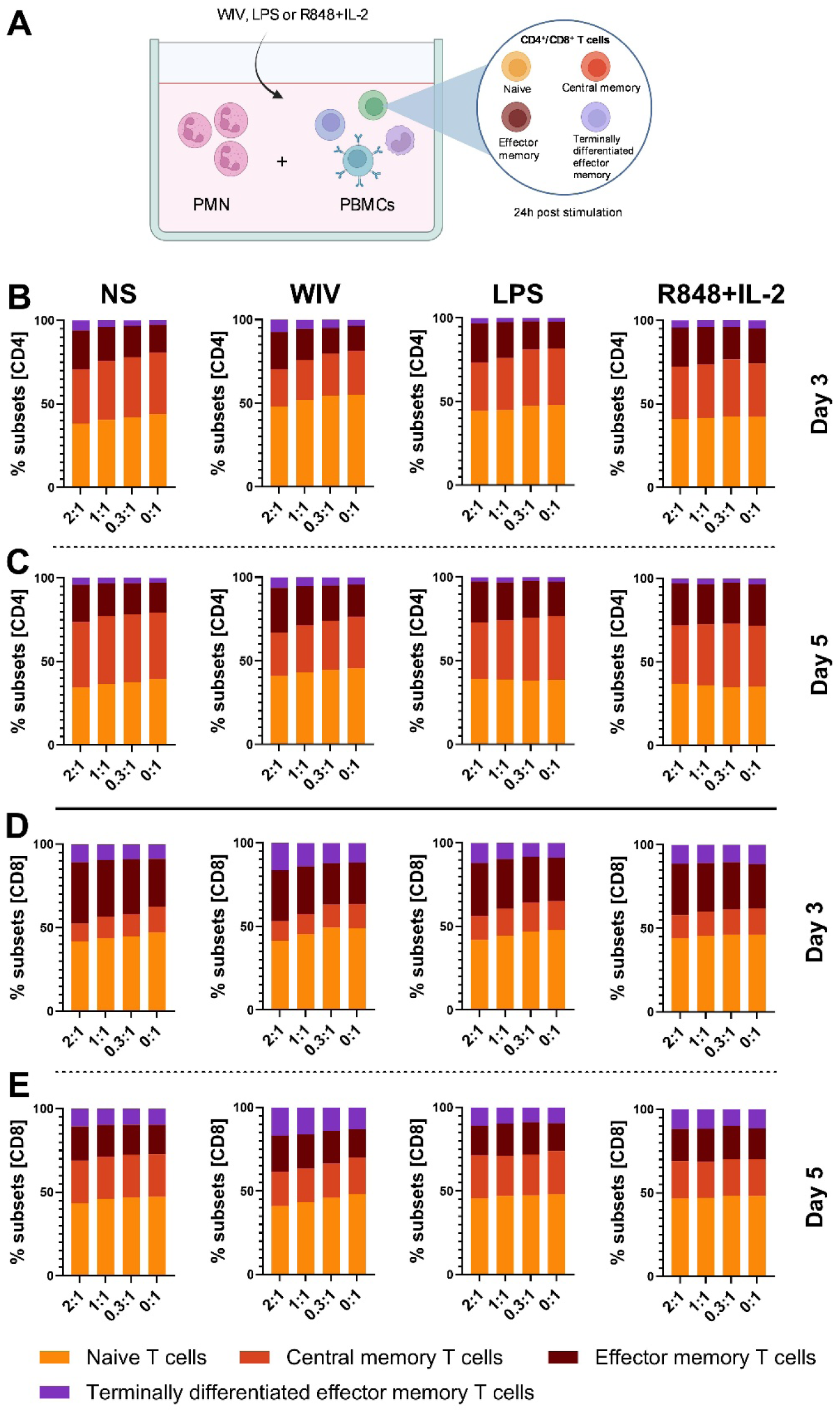
The effect of PMNs on T cell subpopulation frequencies. Autologous PMNs and PBMCs were co-cultured with or without stimulation with WIV or TLR ligands (A). After 3 or 5 days, cells were harvested and T cell subsets were investigated by flow cytometry. The proportions of naïve (T_N_; CCR7^+^CD45^+^), central memory (T_CM_; CCR7^+^CD45^-^), effector memory (T_EM_; CCR7^-^CD45^-^) and terminally differentiated effector memory (T_EMRA_; CCR7^-^CD45^+^) T cells among CD4^+^ (B and C) and CD8^+^ (D and E) T cells were assessed on day 3 (B and D) and day 5 (C and E) using flow cytometry.

T follicular helper (Tfh) cells play a crucial role in enhancing germinal center responses and long-lasting vaccine efficacy [35–37], warranting further investigation. Our study found that, in the absence of stimuli, the proportion of Tfh among CD4^+^ T cells decreased significantly with the presence of PMNs, with the highest PMN:PBMC ratio resulting in the strongest reduction in Tfh (Fig. 6 A, D). This suppressive effect of PMN on Tfh frequencies was most profound on day 3 and became less prominent on day 5 (Fig. 6 A, D). Upon stimulation with WIV, the inhibitory effect of PMNs on Tfh cell frequencies on day 3 was similar as that in non-stimulated cultures (Fig. 6 B). However, on day 5, only a minor decrease in the percentage of Tfh was observed in the presence of WIV (Fig. 6 E) indicating that WIV counteracted PMN-induced suppression of Tfh cells. Notably, the change of Tfh frequency induced by WIV stimulation closely mirrored that seen with R848+IL-2 stimulation (Fig. 6 B, C, E, F), suggesting comparable activation pathways. The frequency of ICOS^+^ Tfh cells in PBMCs increased upon stimulation of the cultures with WIV or LPS. The enhanced activation status of the Tfh cells remained unaffected by the addition of PMNs at all tested ratios, with no significant differences observed on either day 3 or day 5 (Fig. S6 A-F). Thus, Tfh cell activation was not affected by PMNs.

**Fig 6.**
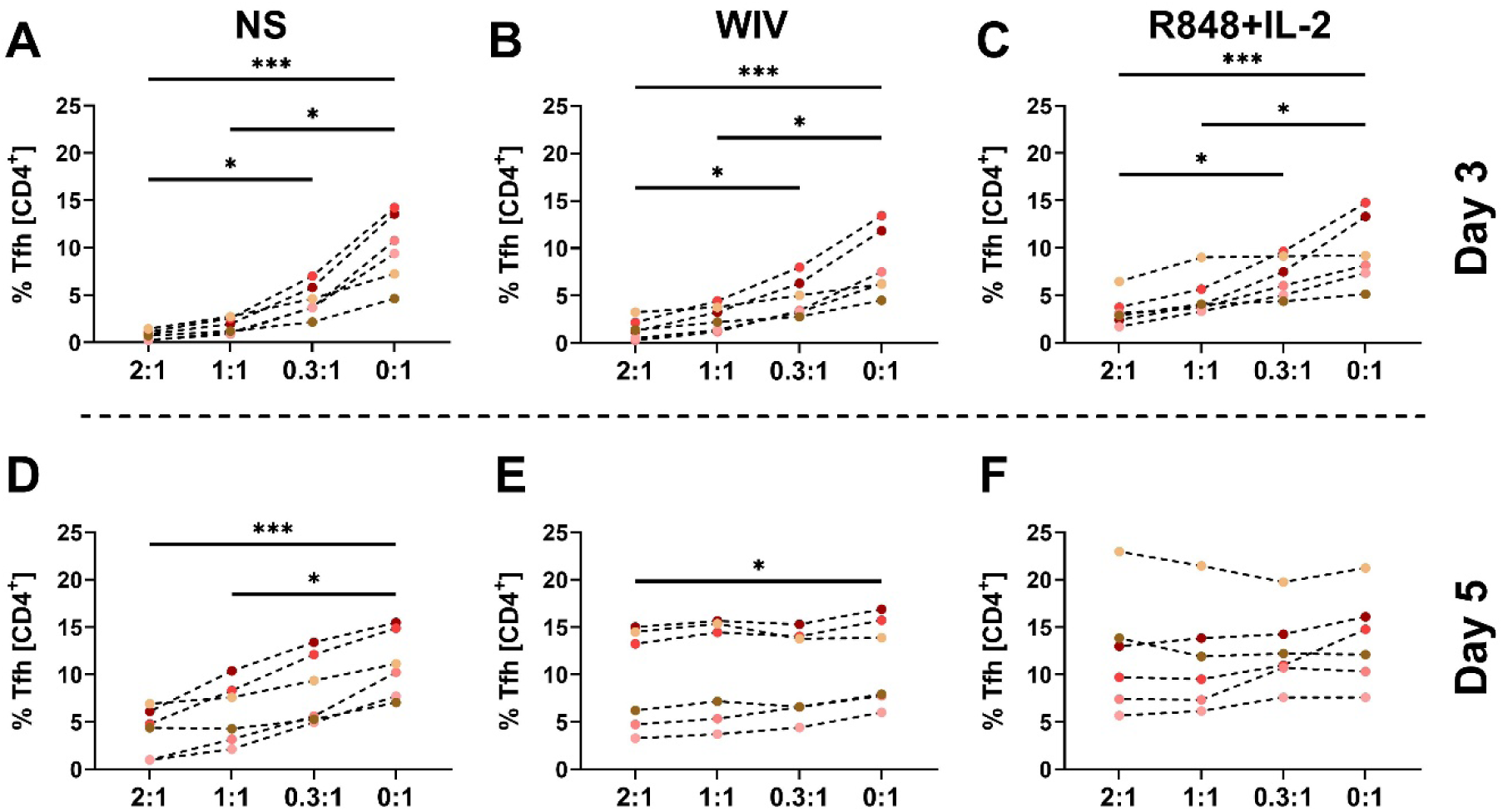
T follicular helper cells in human PMN/PBMC co-cultures. Autologous PMNs and PBMCs were co-cultured at the indicated ratios, either without (A and D) or with stimulation with WIV or R848+IL-2 (B, C, E, and F) for 3 (A-C) or 5 (D-F) days. T follicular helper (Tfh) cells in PBMCs were assessed by flow cytometry. The paired Friedman test was employed to compare all groups, with significance levels denoted as follows: *p < 0.05, and ***p < 0.001.

Taken together, the addition of PMNs slightly promoted the differentiation of naïve T cells to effector subtypes in co-cultured PBMCs but mainly decreased the proportion of Tfh cells unless a strong stimulus like WIV prevented this.

### PMNs alter B cell subsets in WIV-stimulated co-cultures

Next, the changes in B cell subsets in PBMCs co-cultured with different amounts of PMNs were recorded on day 3 and day 5 (Fig. 7 A). After 3 or 5 days of incubation, naïve B cells were the dominant B cell subset in non-stimulated cultures, followed by memory B cells, transitional B cells, and plasmablasts, irrespective of the number of PMNs in the co-culture (Fig. 7B). While stimulation with LPS had only minor effects on this distribution, stimulation with WIV and, to a lesser extent, R848+IL-2 profoundly changed the B cell subset frequencies in PBMCs with strong increases in the fractions of plasmablasts and transitional B cells and decreases in the naïve B cell fraction (Fig. 7 C-E and G-H). The presence of PMNs in the WIV-stimulated cultures further increased the frequency of plasmablasts both after 3 and 5 days of incubation (Fig. 7 C and G). In the R848+IL-2-stimulated cultures, PMNs had only minor effects (Fig. 7 E and I) and in LPS-stimulated cultures PMNs had no effect on plasmablast frequency but increased the frequency of memory B cells and decreased the frequency of transitional B cells (Fig. 7 D and H). These results indicate that PMNs can modify vaccine-imposed effects on B cells.

**Fig 7.**
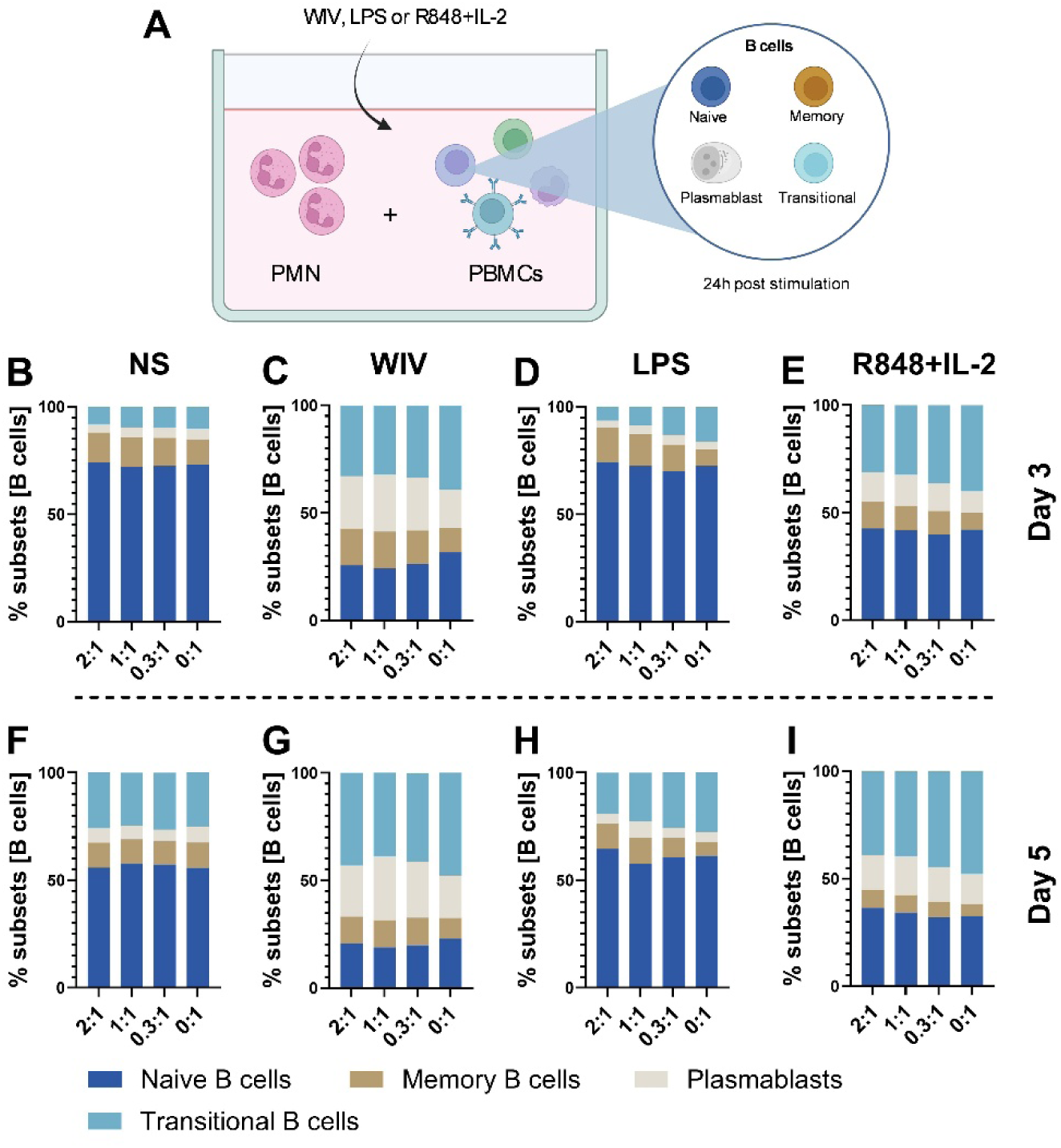
B cell subsets in total B cells of human PMN/PBMC co-cultures. Autologous PMNs and PBMCs were co-cultured at the indicated ratios, either without (B, F) or with stimulation with WIV (C, G), LPS (D, H), or R848+IL-2 (E, I). On day 3 (B-E) and day 5 (F-I), the frequencies of B cell subsets (naïve, transitional, memory B cells and plasmablasts) were measured using flow cytometry.

### PMNs enhance vaccine-induced IgG production in PBMC co-cultures

To investigate the impact of PMNs on humoral immune responses, total IgG levels were measured in supernatants of PMN/PBMC co-cultures harvested after 5 days of stimulation with WIV or TLR ligands. Without stimulation, the total IgG levels in the supernatants of cells cultured at a 2:1 PMN:PBMC ratio were marginally greater than those in cultures without PMNs (0:1) or a small number of PMNs (0.3:1) (Fig. 8A). In the absence of PMNs, stimulation with WIV increased the total IgG levels by a factor of 2 over those in non-stimulated cultures (not shown); the presence of PMNs prompted WIV-stimulated B cells to secrete even higher levels of total IgG than in cultures without PMNs (Fig. 8 B). PMNs also enhanced IgG production in LPS-stimulated cultures (Fig. 8 C) while in R848+IL-2-stimulated, a consistent PMN-induced increase in IgG production was not observed (Fig. 8 D). Overall, the presence of PMNs induced B cells in PBMCs to differentiate into antibody-secreting cells and promoted the WIV-induced production of total IgG.

**Fig 8.**
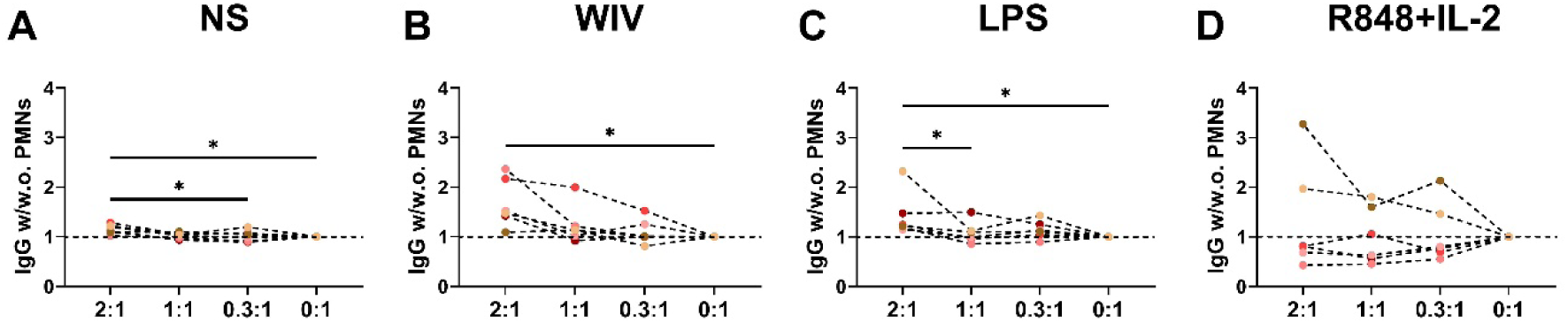
Levels of total IgG in PMN/PBMC co-cultures with and without stimulation. On day 5, cell supernatants from human PMN/PBMC co-cultures were collected and total IgG levels were quantified using ELISA. The fold changes (FC) in total IgG levels for the non-stimulated (A), WIV- (B), LPS- (C), or R848+IL-2- (D) stimulated groups were calculated by dividing the IgG levels of cultures with PMNs (at PMN/PBMC ratios of 2:1, 1:1, and 0.3:1) by those of the corresponding cultures without PMNs (0:1 ratio). A paired Friedman test was performed to assess statistical significance. *p < 0.05.

### PMN addition to PBMC cultures limits vaccine-induced cytokine secretion

In order to understand how the addition of PMNs affects the overall secretion of cytokines, we measured the levels of multiple cytokines in the supernatants of cells with (2:1) or without (0:1) PMN addition after 24 hours of culture. In the absence of stimulation (NS), the addition of PMNs slightly elevated the levels of several cytokines, but statistical significance was not reached (Fig. 9 A-G). Stimulation with WIV increased the levels of MCP-1, CXCL-10, IFNγ, and IL-2 compared to those in non-stimulated cultures, though not significantly (Fig. 9 E, G, J, K). This effect was similar in the absence and presence of PMNs. In contrast, PMN addition to WIV-stimulated PBMCs resulted in a slight reduction of the secretion levels of various other cytokines (Fig. 9 A-C, F, H, I). No to minor effects by PMNs were observed in LPS and R848+IL-2-stimulated conditions. Taken together, PMNs had no or a slightly suppressive effect on the production of cytokines induced by WIV, LPS, or R848+IL-2.

**Fig. 9.**
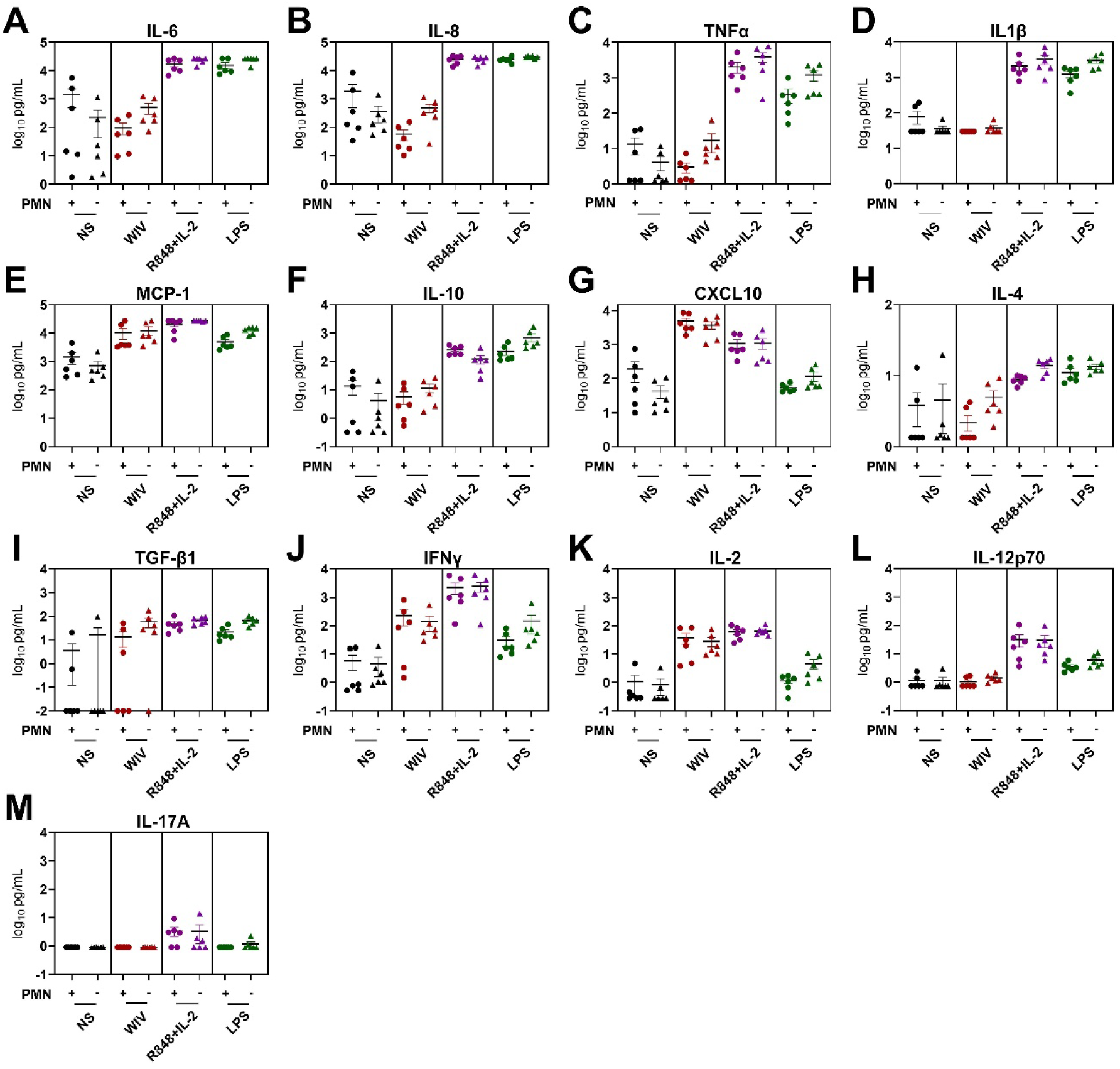
Quantification of cytokine levels using LEGENDplex™ multiplex assay. After 24 hours of treatment with WIV, R848+IL-2, or LPS, supernatants from cell cultures with PMN addition (2:1) and without PMN addition (0:1) were collected. Levels of multiple cytokines in the harvested supernatants, including IL-6 (A), IL-8 (B), TNF-α (C), IL-1β (D), MCP-1 (E), IL-10 (F), CXCL10 (G), IL-4 (H), TGF-β1 (I), IFN-γ (J), IL-2 (K), IL-12p70 (L), and IL-17A (M), were quantified using bead-based flow cytometry. The horizontal lines represent the mean ± SEM of 6 independent donors.

## Discussion

Our study aimed to investigate the response of PMNs to vaccines and to explore the potential interactions between PMNs and other immune cells using an *in vitro* (co-)culture system. Our findings demonstrate that PMNs when cultured alone show minimal responses to a WIV influenza vaccine and thus suggest they require additional stimulants for full activation. In PMN/PBMC co-cultures, PMNs modulated WIV-induced immune cell responses by: (1) altering monocyte subsets, (2) enhancing myeloid cell activation, (3) suppressing Tfh cells (unless WIV was present), and (4) driving antibody production by B cells. These effects occurred without significant cytokine changes, suggesting alternative mechanisms.

As for the monocultures of PMNs in this study, interestingly, WIV slightly extended the survival of PMNs and sustained surface CD16 expression on PMNs compared to unstimulated PMNs. These results align with earlier findings showing a link between apoptosis of PMNs and a reduction in CD16 expression [38–40]. Thus, vaccines can modulate PMN characteristics such as survival and expression of immune receptors which may change the way the cells contribute to vaccine-induced immune activation.

Regarding the impact of PMNs on innate immune cells, the addition of PMNs to PBMCs induced the expression of HLA-DR and CD86 on the surface of CD14^-^CD11c^+^ cells, a cell type consisting mainly of conventional DCs (Fig. 4), which is consistent with previous studies [41]. The activation of DCs by PMNs may involve the physical interaction of the C-type lectin DC-SIGN on the surface of DCs and Mac-1 on PMNs [42]. DC activation by PMNs may also be triggered by PMN-derived TNFα [43]. However, we observed only slightly increased levels of TNFα in the cell supernatants of PMN/PBMC co-cultures (Fig. 9C). Moreover, we noted that, in the presence of WIV, PMNs impacted on the relative frequencies of monocyte subsets, decreasing the frequency of CMs (CD14^hi^/CD16^-^) and increasing the frequencies of IMs (CD14^hi^/CD16^+^) and NCMs (CD14^+^/CD16^+^). IMs exhibit superior antigen-presenting capabilities and MHC II expression, essential for T cell activation, while CMs have a higher capacity for cytokine production, phagocytosis, and adhesion [44–46]. This suggests that under vaccine stimulation, PMNs could drive monocytes to differentiate towards antigen-presenting cells. However, the specific mechanism of how PMNs promote monocyte differentiation and whether IM promotion is a general phenomenon or happens only under certain conditions is still unclear and requires further investigation.

In recent years, researchers have studied the effects of PMNs on adaptive immune cells both *in vitro* and *in vivo*. Current findings suggest that PMNs may either enhance or suppress T cell activity, the impact on T cells likely depending on the activation state of the PMNs [47, 48]. The mechanisms by which PMNs affect T cells are complicated. PMNs can directly present antigens to T cells, interact with dendritic cells or macrophages to assist in antigen presentation to T cells, produce neutrophil extracellular traps (NETs) and myeloperoxidase (MPO), or secrete various cytokines or chemokines to modulate T cell activity [25, 41, 42, 49–56]. In our study, T cells in PBMCs co-cultured with PMNs exhibited enhanced differentiation to T_EM_ and T_EMRA,_ while at the same time the frequency of Tfh declined. Considering that, without stimulation, none of the cytokines we determined showed a significant change upon the addition of PMNs, we speculate that the influence of PMNs on T cells in PBMCs may largely stem from direct physical contact and/or by so far unrevealed factors affecting the T cells or other immune cells. Co-culturing PMNs and T cells and adding in other purified immune cells and/or using Transwell technology to separate specific cell populations might help to elucidate the mechanisms of PMN effects on T cells.

As for humoral immune responses, our results indicate that the presence of PMNs indeed promoted the differentiation and antibody production capacity of B cells under WIV stimulation. This is consistent with the basic conclusions of other studies [57–59]. In the absence of vaccine stimulation, PMNs did not affect the differentiation of B cells to antibody-secreting cells. This might be related to the reduction in Tfh numbers in non-stimulated PMN/PBMC co-cultures. Tfh cells assist B cells and are *in vivo* crucial for the formation of germinal centers, and the development of memory B cells. Moreover, they promote the differentiation of B cells to antibody-producing plasmablasts and plasma cells [60]. While with increasing numbers of PMNs Tfh frequencies declined in non-stimulated co-cultures, in the presence of WIV, Tfh numbers did not decrease when PMNs were present, concomitant with increased antibody production. Furthermore, under WIV stimulation in the presence of PMNs, the levels of CXCL10 and IFN-γ in the cell supernatants either increased or remained slightly elevated (Fig. 8 G, J, K). CXCL10 plays a role in the migration and activation of Tfh cells [61]. IFN-γ can affect Tfh cell differentiation and support specific aspects of B cell differentiation [62, 63]. Thus, the generation of antibody-secreting cells and the production of antibodies might be affected by effects of PMNs on Tfh cells. However, formal proof of a causal role of PMNs in these processes still needs to be provided.

It is generally accepted that PMNs play an important role in inflammation. In our experiments, we observed that in the absence of stimulation, PMNs enhanced the release of inflammatory cytokines and chemokines. However, under stimulation with different vaccines or TLR ligands, the presence of PMNs led to a decrease in most cytokines. Gresnigt et al. similarly observed in co-culture experiments involving PMNs and PBMCs that the presence of PMNs reduced LPS-induced IL-1β and TNF-α production, aligning with our findings [64]. Such anti-inflammatory effects of PMNs may result from the release of soluble substances, such as proteases and/or anti-inflammatory factors, the scavenging of inflammatory factors by apoptotic PMNs, production of microparticles like microvesicles and ectosomes, and physical contact [64–66]. Several recent *in vivo* studies delved into the role of PMNs in vaccination, considering factors such as vaccine types, vectors, and adjuvants, and confirmed a role of PMNs in the transport of antigen to lymph nodes and the activation of T cell immunity [8, 9, 58, 67]. However, these studies were primarily conducted in mice. Given the significant differences between murine and human PMNs, it is essential to use human PMNs for research. *In vitro* studies using human PMNs co-cultured with other immune cells at specific ratios typically demonstrate responses to TLR ligands, heat-killed *Candida albicans,* tetanus toxoid, cytomegalovirus (CMV) pp65, or influenza hemagglutinin (HA) [25, 29, 41, 48, 53, 64, 68, 69]. Moreover, most of these *in vitro* studies focused on interactions between PMNs and a single type of immune cell. Our study incorporated different stimuli into a relatively complex co-culture of human PMNs and PBMCs to conduct a comprehensive investigation of immune cells, from phenotype to activation to cytokine/chemokine release.

Our study has some limitations. Given that previous studies indicated that the impact of PMNs on innate immune cells and T cells likely requires physical contact, we decided in our experiment to co-culture PMNs directly with PBMCs. However, this setup did not allow us to discriminate between physical contact and soluble factors as being most important in cell activation. Future experiments should make use of devices like Transwell® plates that allow the culture of PBMCs and PMNs in separate compartments or the removal of PMNs at specific times. Furthermore, there are concerns that the continuous accumulation of metabolic byproducts from rapidly dying PMNs in co-culture might stress other immune cells. Conducting further and more comprehensive observations of PMN effects on B and T cells is therefore challenging. However, optimizing the culture environment might foster a breakthrough to solve this problem.

Taken together, our study demonstrates bi-directional effects exhibited by PMNs and WIV. WIV delayed PMN apoptosis in pure PMN cultures. In PMN/PBMC co-cultures, PMNs modulated WIV-induced monocyte changes and enhanced CD14⁻CD11c⁺ cell activation. They promoted effector T cell differentiation and boosted B cell antibody production, including WIV-induced IgG. PMNs increased baseline inflammation but suppressed vaccine-triggered cytokine release, possibly demonstrating context-dependent immunomodulation that balances pro-inflammatory and adaptive immune activation. These results point to a modulating role of PMNs in vaccine-induced immune responses and imply that PMNs should be taken into account during vaccine development and assessment.

